# Glucagon-like peptide-1 receptor agonist, semaglutide, attenuates intravenous self-administration of fentanyl in female rats

**DOI:** 10.64898/2026.05.19.726324

**Authors:** Kaitlyn E. Rojas, Sarah C. Gee, Chris L. Wernette, Eric X. Wang, Emily T. Nguyen, Jacques D. Nguyen

## Abstract

Current treatments for opioid use disorder (OUD) have major barriers to access. As such, researching new potential therapies for OUD is important to public health. Previous research has implicated glucagon-like peptide-1 (GLP-1) receptor agonists in decreasing the use of addictive substances by animals. In this study, female Wistar rats (N=32) were surgically implanted with jugular catheters and trained to self-administer fentanyl at a fixed-ratio 1 (FR1) schedule of reinforcement for 21 sessions under short- (ShA; 1 hour) or long-access (LgA; 8 hours) conditions. Next, the animals received injections of semaglutide (0.1 mg/kg, s.c.) or saline (0.9% NaCl, s.c.) prior to another FR1 session. The animals underwent a progressive ratio (PR) schedule of reinforcement while receiving saline (i.v.) or fentanyl (0.625-10 µg/kg/inf, i.v.) and semaglutide (0.1 mg/kg, s.c.) or saline (s.c.). Next, the animals underwent a semaglutide (0-0.1 mg/kg, s.c.) dose response procedure at FR1 and a single dose of fentanyl (2.5 µg/kg/inf, i.v.). Following drug discontinuation, spontaneous locomotor activity and withdrawal-like symptoms were measured. Semaglutide dose-dependently decreased fentanyl rewards under ShA and LgA conditions (p<0.05). Under a PR, semaglutide significantly decreased breakpoint (p<0.05), suggesting semaglutide decreases motivation to self-administer fentanyl. Semaglutide-treated ShA animals displayed significantly less withdrawal-like behavior (p<0.05) but not LgA animals. Overall, these findings suggest semaglutide may modulate motivation to seek opioid reward and could be useful in the development of pharmacotherapies to address OUD.

## 1. INTRODUCTION

The recent decade has seen a surge of synthetic opioid related deaths [1]. The U.S. Drug Enforcement Agency stated that, in 2023 alone, they seized over 79 million illicit pills containing fentanyl [2]. Due to the rise in fentanyl-laced pills, treatment for opioid use disorder (OUD), defined as “a problematic pattern of opioid use leading to clinically significant impairment or distress” [3], has been of high priority [4]. In 2022, an estimated 9.4 million U.S. adults required treatment for OUD [5], and current treatments for OUD are underused and retain major barriers to patient access despite their ability to greatly reduce mortality from all causes [6]. Approximately 38,409 people in the U.S. died from a synthetic opioid overdose from September 2024 to September 2025 [7]. With this knowledge in mind, researching better potential treatments for OUD is of the utmost importance to public health.

Glucagon-like peptide-1 (GLP-1) is a hormone synthesized and released from the enteroendocrine cells of the small intestine and the nucleus of the solitary tract of the brainstem in response to food consumption and it exerts its effects by binding to endogenous GLP-1 receptors [8-10]. Activation of GLP-1 receptors increases insulin secretion, decreases blood sugar and glucagon release, decreases gastric motility, and increases satiety [11-14]. The ability of GLP-1 to allow people to feel full for longer has garnered interest in exogenous GLP-1 receptor agonists, i.e. semaglutide (Ozempic®, Wegovy®), liraglutide, exendin-4, tirzepatide, etc., as weight-loss medications, while the ability of GLP-1 receptor agonists (GLP-1RAs) to decrease blood sugar and to increase insulin secretion have led to their use in treating Type 2 Diabetes [15]. Between 2011 to 2023, 871,854 individuals were prescribed a GLP-1RA [16].

The success of GLP-1RAs in decreasing motivation for food rewards sparked curiosity in whether this decrease in motivation could be applied to substance use disorder. This has recently gained media attention with *Nature* [17] and *Science* [18] publishing news articles highlighting the ongoing research in this growing field. Previous studies found that GLP-1RAs decreased alcohol use both in animal models [19-21] and in patients with alcohol use disorder [22]. Other studies found that GLP-1RAs and GLP-1 signaling reduced cocaine self-administration in animals and in patients with cocaine use disorder [23-27]. Similarly, several studies demonstrated the ability of GLP-1RAs to decrease self-administration of opioids, such as heroin [28-31], oxycodone [32], and fentanyl [33-36], in rodent models of OUD. Presently, no studies have examined the effects of semaglutide, a well-known GLP-1RA, on self-administration of opioid drugs. Additionally, most studies have focused on the impacts of GLP-1RAs on *reinstatement* of drug-seeking and -taking [30-32,35-38], not on expression of *acquisition* or *escalation* of self-administration behavior. Lastly, the vast majority of studies examined the effects of GLP-1 receptor agonists on drug self-administration in male animals only. The present study determined the effects of semaglutide on self-administration of intravenous fentanyl in female Wistar rats.

## 2. METHODS AND MATERIALS

### 2.1 Animals

Female Wistar rats (N=32) underwent jugular catheterization and arrived at Baylor University at 10-11 weeks of age (Envigo; Indianapolis, IN). Animals were pair-housed in a temperature-controlled (23ºC±1ºC) vivarium on a 12:12h reverse light cycle and given *ad libitum* access to food (5V6R PicoLab® Select Rodent 50 IF/6F) and water. All experiments occurred during the animals’ scotophase. Daily, the subjects were handled, weighed, and had their catheters flushed with a solution of cefazolin in heparinized saline. All aspects of this study were approved by the Baylor University Institutional Animal Care and Use Committee and performed in accordance with their guidelines.

### 2.2 Drugs

Fentanyl citrate powder (Millipore Sigma; St. Louis, MO) was dissolved in 0.9% sodium chloride (NaCl) sterile saline to produce the solutions used for fentanyl self-administration. Each infusion (0.1 mL) provided an intravenous dose of 0.625, 2.5, or 10 µg/kg/inf, adjusting based on the average body weight of the subjects. All solutions were prepared weekly. Fentanyl dosing was based off of previous self-administration work in our lab and others [34,39,40]. Semaglutide (Cayman Chemical Company; Ann Arbor, MI) was dissolved in 0.9% NaCl sterile saline to produce a stock solution of 0.1 mg/mL. The stock solution was further diluted using sterile saline to prepare the lower concentration solutions: 0.01 and 0.03 mg/kg. Semaglutide dosing was consistent with previous studies [19,20,41]. Cefazolin (1 gram; WG Critical Care LLC; Paramus, NJ) was dissolved in 12.5 mL of heparinized saline. Animals received approximately 0.2-0.3 mL (i.v.) of this solution daily. Heparinized saline was prepared by dispensing 1 mL of heparin (1,000 USP/mL; Sagent Pharmaceuticals; Schaumburg, IL) into 30 mL bacteriostatic saline (0.9% sodium chloride; Hospira, Inc.; Lake Forest, IL).

### 2.3 Intravenous Self-Administration

Throughout intravenous self-administration experiments, animals were placed in operant chambers within sound attenuating boxes (MedAssociates; St. Albans, VT). These chambers were equipped with a cue light and two levers: an active lever, pressing which would result in an infusion of saline (0.9% NaCl, i.v.) or fentanyl (0.625-10 µg/kg/inf, i.v.) depending on the session, and an inactive lever, pressing which resulted in no consequence. Prior to all self-administration sessions, experimenters flushed the catheters of the animals with a solution of Cefazolin in heparinized saline, prepared following the procedure described in the previous section. The catheters of the animals were also flushed with this solution all other days to ensure proper catheter maintenance. Following program initiation at the start of each intravenous self-administration session, animals received a 2-second infusion to purge any remaining catheter-flushing solution. This ensured that the first correct response would warrant the correct dose of fentanyl rather than be diluted by the cefazolin in heparinized saline solution. After completion of each session, the subjects’ catheters were flushed with a solution of heparinized saline before return to their home cages.

Fixed-ratio 1 (FR1) sessions were structured such that sessions ended after 1 hour (short-access; ShA) or 8 hours (long-access; LgA) or after an animal had earned the maximum of 300 infusions, whichever occurred first. Additionally, under FR1 animals received a priming infusion of fentanyl if no responses were observed within 30 minutes of the session start. Between ShA and LgA sessions, the operant chambers were cleaned with a solution of 30% isopropanol. Animals were trained to self-administer intravenous fentanyl (2.5 µg/kg/inf) at an FR1 schedule of reinforcement for 21 sessions under either ShA (1 hour) or LgA (8 hour) conditions. On the 22nd session, animals continued this FR1 schedule and received an injection of either semaglutide (0.1 mg/kg, s.c.) or vehicle (0.9% saline, s.c.) 30 minutes prior to their session start.

Then, the animals proceeded to a PR schedule of reinforcement where they were tested across 4 sessions at variable doses of fentanyl (0-10 µg/kg/inf, i.v.) in a randomized order while receiving a stable dose of semaglutide (0.1 mg/kg, s.c.) or vehicle (0.9% saline, s.c.). Progressive ratio (PR) sessions were structured such that sessions ended after 300 infusions were earned, 180 minutes, or the animal reached breakpoint, whichever occurred first. The animal was defined as reaching breakpoint after not responding for 30 minutes after receiving their last infusion. Breakpoint values indicate the number of correct lever responses necessary to receive an infusion at which the animals stopped responding.

Lastly, rats received an approximately 60-hour abstinence period before beginning a semaglutide dose-response procedure. For this dose response, semaglutide-experienced animals remained at a stable dose of fentanyl (2.5 µg/kg/inf, i.v.) under the original FR1 schedule of reinforcement and received variable doses of semaglutide (0-0.1 mg/kg, s.c.) in ascending order across 4 sessions, while the vehicle-experienced animals self-administered fentanyl under FR1.

### 2.4 Locomotor Activity Assessment

To determine whether semaglutide administration would alter spontaneous locomotor activity, all animals underwent a locomotor activity assessment 16-22 hours following fentanyl discontinuation. Animals received injections of either semaglutide (0.1 mg/kg, s.c.) or vehicle (0.9% saline, s.c.) 30 minutes prior to entering Superflex Open Field boxes (Omnitech Electronics, Inc.; Columbus, OH) for 30 minutes. Boxes were cleaned with deionized water and dried between subjects. Fusion v6.5 SuperFlex Edition (Omnitech Electronics, Inc.; Columbus, OH) recorded the laser beams broken by the movement of the animals and provided data on ambulatory activity count, horizontal activity count, vertical activity count, total distance travelled, and rest time. This data was then exported for statistical analyses.

The software measured ambulatory activity by ambulatory activity count, defined as the number of beam breaks while the animal is ambulating, which does not include stereotypy behavior. Horizontal activity was measured by horizontal activity count, defined as the number of horizontal beam breaks. Vertical activity was measured by a vertical activity count, defined as the cumulative vertical beam breaks. Total distance (cm) was the total distance that the animal traveled, and the animal’s location was defined by the centroid model, focusing on the center of mass of the subject. Rest time (s) was defined as the length of time that the animal spent at rest, where the subject did not break any beams for 1 second or more. A resting period was defined as a period of inactivity greater than or equal to 1 second. The Open Field SuperFlex boxes were partitioned into center and perimeter areas and evaluated as an open field test. Open field is a widely used measure of anxiety-like behavior in rodent models [42]. Here, we compared duration of time spent in the center or perimeter, using the centroid model, which considers the animal to be within a zone (i.e. in the center or perimeter) if the center of the subject is within the zone.

### 2.5 Withdrawal Scoring

Following locomotor activity assessment, the animals were recorded for scoring to assess signs of opioid withdrawal-like symptoms. These recordings lasted 30 minutes and occurred approximately 18-24 hours following fentanyl discontinuation. The recordings were scored by 3 treatment-blinded researchers. The average of these scores was used in statistical analyses. Scorers evaluated animals for abdominal constrictions, wet dog shakes, facial fasciculations, ptosis, abnormal posture, and genital grooming, based on a scale adapted from Geller and Holtzman [43]. Abdominal constrictions and wet dog shakes were scored as graded symptoms, while facial fasciculations, ptosis, abnormal posture, and genital grooming were scored as “all-or-nothing” symptoms. Table 1 further details the weight of each symptom.

**Table 1.** Point system used for opioid withdrawal scoring, adapted from Geller and Holtzman (1978). Abdominal constrictions and wet dog shakes were scored as weighted symptoms where values varied based on number of occurrences. Facial fasciculations/teeth chatter, ptosis, abnormal posture, and genital grooming were considered fixed symptoms, with no difference in scoring based on the number of occurrences. If any symptoms were not observed during the recording, the score was recorded as zero.

| Number of occurrences | Abdominal Constrictions | Wet Dog Shakes | Facial Fasciculations/Teeth Chatter | Ptosis | Abnormal Posture | Genital Grooming |
| --- | --- | --- | --- | --- | --- | --- |
| 1 | 2 points | 2 points | 2 points | 2 points | 2 points | 2 points |
| 2 | 4 points | 2 points | 2 points | 2 points | 2 points | 2 points |
| ≥3 | 6 points | 4 points | 2 points | 2 points | 2 points | 2 points |

### 2.6 Statistical Analysis

All analyses were performed using GraphPad Prism (v10.6.1; Boston, MA). Data were analyzed using Welch’s t-test, One-Way ANOVA, Two-Way ANOVA, or Three-Way ANOVA, as appropriate. The main effects assessed included Treatment, Dose, Access Duration, and/or Session. Tukey’s post-hoc analysis for multiple comparisons was used given a significant main effect was confirmed. Results were considered significant for p<0.05. All data are represented as mean ±SEM.

## 3. RESULTS

### 3.1 Female Wistar rats escalated drug-taking for fentanyl under LgA but not ShA conditions

A Two-Way ANOVA and a Tukey’s post-hoc analysis for multiple comparisons were used to examine changes in fentanyl self-administration throughout acquisition. Analyses confirmed effects of Session (F(20,600)=7.868; p<0.05), Access Duration (i.e. ShA vs LgA; F(1,30)=100.4; p<0.05), and Subject (F(30,600)=14.40; p<0.05). Session x Access Duration (F(20,600)=5.456; p<0.05). As shown in **Figure 1A**, Tukey’s post-hoc analysis confirmed that animals trained under LgA conditions began earning significantly more rewards, or infusions of drug, compared to their first session from session 3 and onward (p<0.05). However, there were no sessions in which the ShA animals received significantly more rewards compared to the first session.

**Figure 1.**
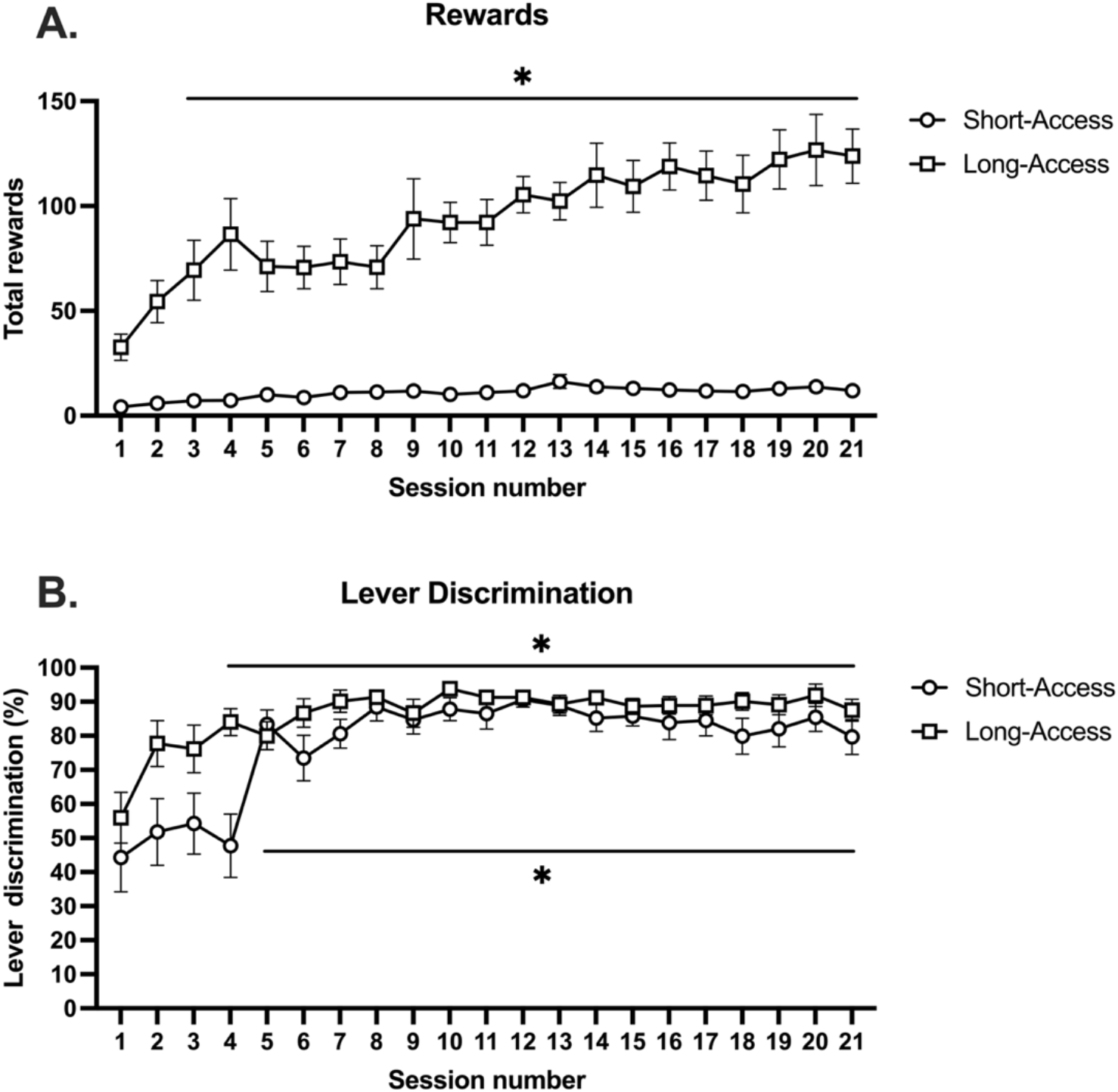
**(A)** Total rewards (mean ± SEM) earned by ShA and LgA animals during acquisition. * indicates a significant increase (p<0.05) in lever pressing for animals trained under long access conditions compared to session 1. **(B)** Percent lever discrimination (mean ± SEM) for ShA and LgA animals during acquisition. * indicates a significant increase in lever discrimination from session 5 onward compared to session 1 (p<0.05) for ShA animals and from session 4 onward compared to session 1 (p<0.05) for LgA animals.

### 3.2 Female Wistar rats trained under LgA conditions significantly increased lever discrimination earlier than those trained under ShA conditions

A Two-Way ANOVA and Tukey’s post-hoc analysis for multiple comparisons were used to examine changes in lever discrimination during acquisition. Analyses confirmed effects of Session (F(20,600)=11.97; p<0.05), Access Duration (F(1,30)=7.923; p<0.05), Subject (F(30,600)=5.008; p<0.05), and Session x Access Duration (F(20,600)=2.175; p<0.05). As shown in **Figure 1B**, Tukey’s post-hoc analysis found that ShA animals self-administered significantly more than the first session from session 5 onward (p<0.05), while LgA animals self-administered significantly more than they did in the first session from session 4 onward (p<0.05).

### 3.3 Effects of semaglutide on expression of fentanyl self-administration under ShA conditions

Following 21 sessions at a FR1 schedule of reinforcement, ShA animals received an injection of either saline (0.9%, s.c.) or semaglutide (0.1 mg/kg, s.c.) 30 minutes prior to the start of their session. As shown in **Figure 2A**, a two-tailed Welch’s t-test confirmed that animals receiving semaglutide injections received significantly less fentanyl rewards than animals receiving saline (p<0.05). For this same session, another two-tailed Welch’s t-test revealed no significant difference in lever discrimination between animals receiving saline versus semaglutide shown in **Figure 2B**.

**Figure 2.**
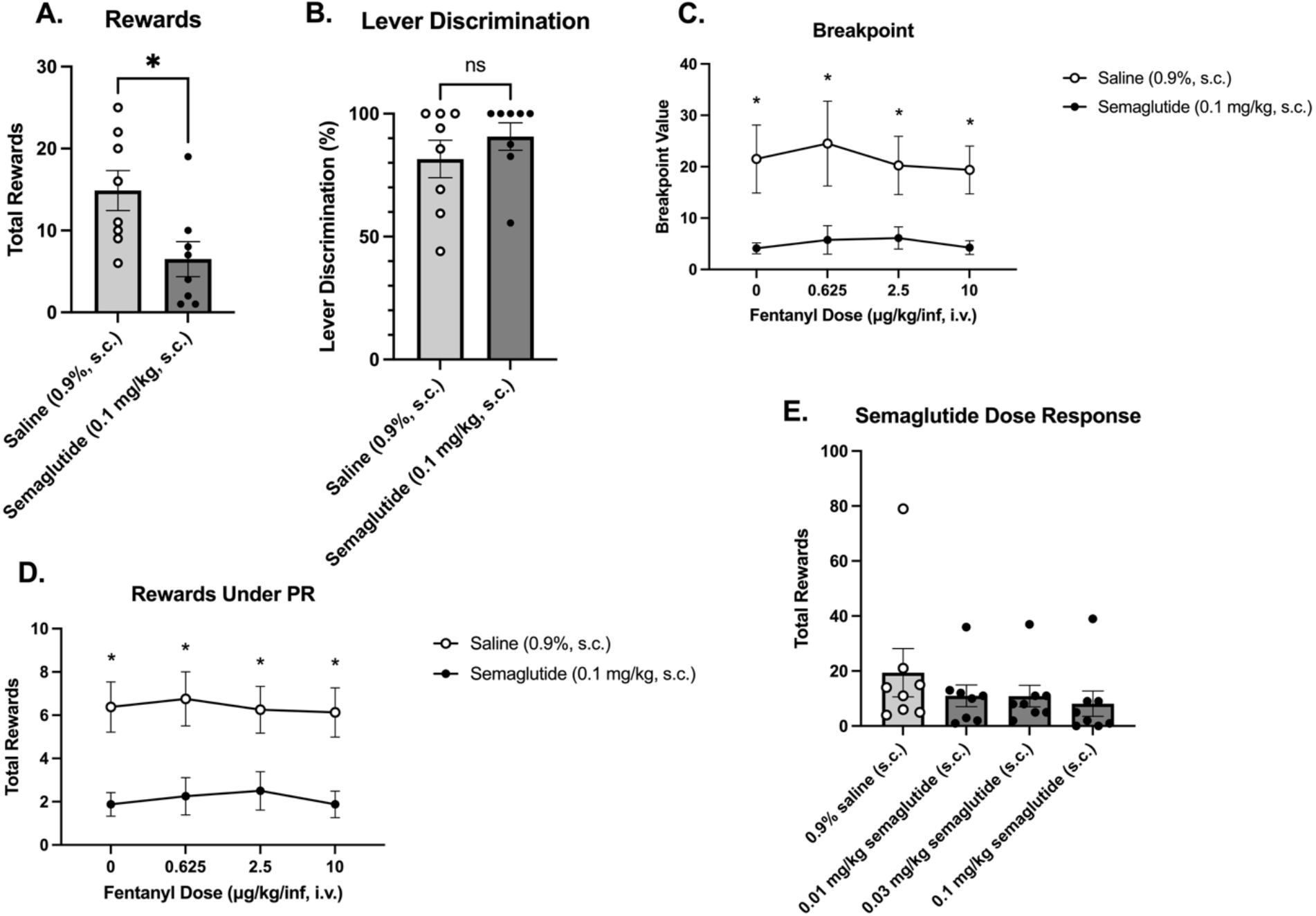
**(A)** Mean rewards earned (±SEM) by ShA animals under FR1 following injections of saline or semaglutide. * indicates a significant decrease in rewards earned by animals receiving semaglutide versus saline (p<0.05). **(B)** Mean percent lever discrimination (±SEM) for ShA animals following injections of saline or semaglutide. No significant effects were observed. **(C)** Mean breakpoint values (±SEM) for ShA animals receiving saline or semaglutide across multiple doses of saline or fentanyl under a PR schedule of reinforcement. * indicates a significant decrease in breakpoint for animals receiving semaglutide as compared to saline for each dose of saline or fentanyl (p<0.05). **(D)** Mean rewards earned during PR (±SEM) by ShA animals receiving injections of saline versus semaglutide. * indicates a significant decrease in rewards earned for animals receiving semaglutide as compared to saline across multiple doses of saline or fentanyl. **(E)** Mean rewards earned (±SEM) by ShA animals under FR1 across escalating doses of semaglutide. No significant effects were observed.

After the 22 total FR1 sessions, ShA animals proceeded to 4 sessions at a PR schedule of reinforcement where they received injections of either saline (0.9%, s.c.) or semaglutide (0.1 mg/kg, s.c.) 30 minutes prior to their session start and where they self-administered saline or fentanyl (0.625-10 µg/kg/inf, i.v.) in a randomized order. A Two-Way ANOVA of breakpoint across fentanyl doses confirmed a significant main effect of Treatment (F(1,56)=23.63; p<0.05). As shown in **Figure 2C**, Tukey’s post-hoc analysis revealed significant differences in breakpoint for animals receiving saline versus semaglutide for animals receiving saline and at all tested doses of fentanyl (p<0.05). Furthermore, a Two-Way ANOVA of rewards earned during PR confirmed a significant main effect of Treatment (F(1,56)=38.08; p<0.05). Tukey’s post-hoc analysis revealed a significant decrease in rewards earned at all tested doses of saline or fentanyl for animals receiving semaglutide versus saline (p<0.05; **Figure 2D**).

Following the PR sessions and an approximately 70-hour abstinence period where animals did not receive fentanyl or semaglutide, the semaglutide-experienced ShA animals completed a dose-response assessment while receiving a 2.5 µg/kg/inf (i.v.) dose of fentanyl and escalating doses of saline (0.9%, s.c.) and semaglutide (0.01-0.1 mg/kg, s.c.). While animals receiving semaglutide self-administered less fentanyl infusions than those receiving saline, an ordinary One-Way ANOVA revealed no significant differences (**Figure 2E**).

### 3.4 Effects of semaglutide on expression of fentanyl self-administration under LgA conditions

After 21 FR1 sessions, LgA animals completed another FR1 session at a 2.5 µg/kg/inf (i.v.) dose of fentanyl and received either saline (0.9%, s.c.) or semaglutide (0.1 mg/kg, s.c.) 30 minutes prior to their session start. A two-tailed Welch’s t-test showed that animals injected with semaglutide injections self-administered significantly less fentanyl infusions than animals injected with saline (p<0.05; **Figure 3A**). Another two-tailed Welch’s t-test revealed that there was no significant difference in lever discrimination for animals receiving semaglutide as compared to those receiving saline (p<0.05; **Figure 3B**).

**Figure 3.**
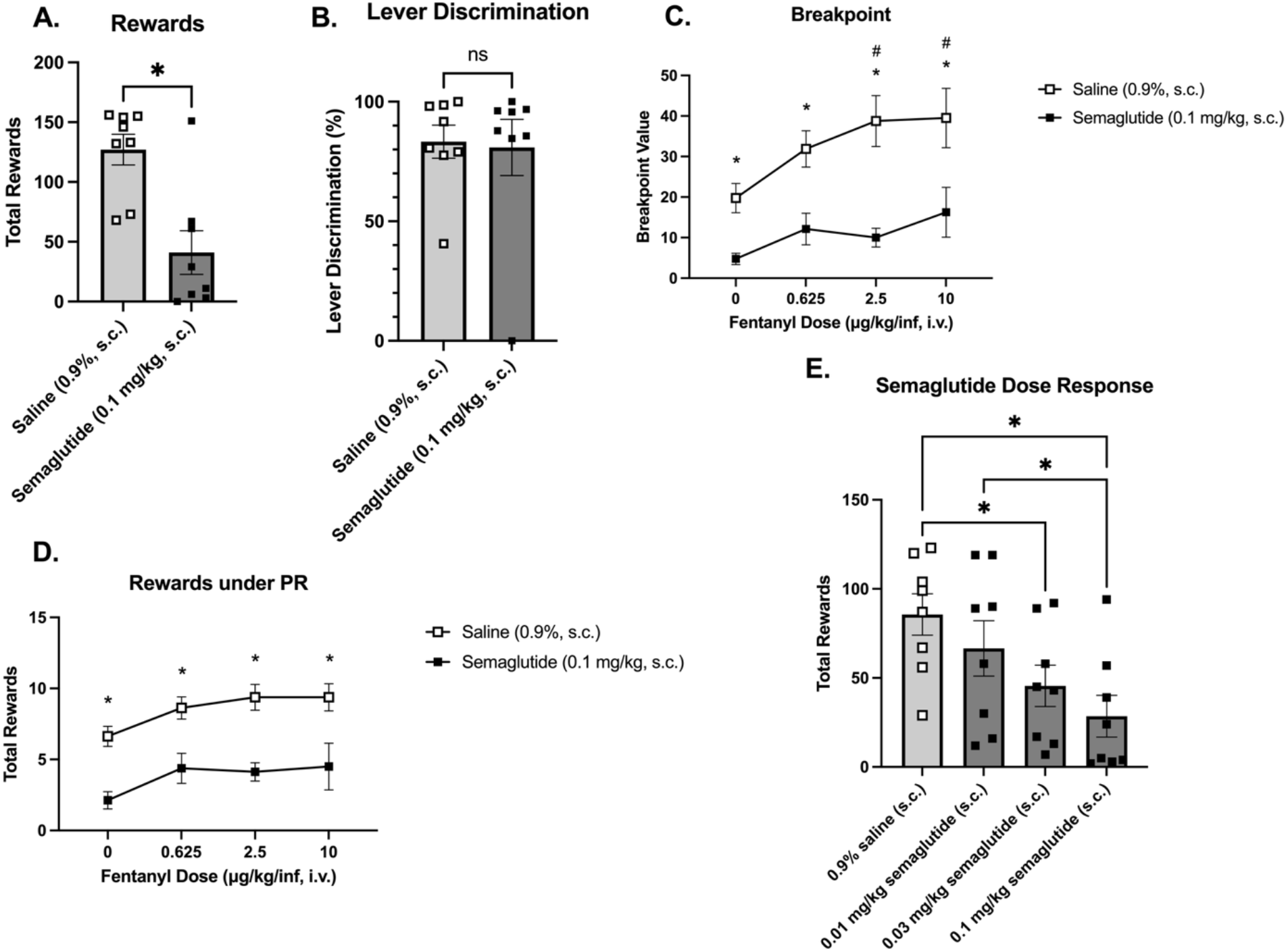
**(A)** Mean rewards earned (±SEM) by LgA animals under FR1 following injections of saline or semaglutide. * indicates a significant decrease in rewards earned by animals receiving semaglutide versus saline (p<0.05). **(B)** Mean percent lever discrimination (±SEM) for LgA animals following injections of saline or semaglutide. No significant effects were observed. **(C)** Mean breakpoint values (±SEM) for LgA animals receiving saline or semaglutide across multiple doses of saline or fentanyl under a PR schedule of reinforcement. * indicates a significant decrease in breakpoint for animals receiving semaglutide as compared to saline for each dose of saline or fentanyl (p<0.05). # indicates a significant difference in rewards earned by saline animals at the 2.5 and 10 µg/kg/inf dose of fentanyl (i.v.) compared to those self-administering saline (0.9%, i.v.; p<0.05). **(D)** Mean rewards earned during PR (±SEM) by LgA animals receiving injections of saline versus semaglutide. * indicates a significant decrease in rewards earned for animals receiving semaglutide as compared to saline across multiple doses of saline or fentanyl. **(E)** Mean rewards earned (±SEM) by LgA animals under FR1 across escalating doses of semaglutide. * indicates a significant difference in rewards earned by animals between doses of saline or semaglutide (p<0.05).

After a total of 22 FR1 sessions, the LgA animals completed a PR schedule of reinforcement where they self-administered saline or fentanyl (0.625-10 µg/kg/inf, i.v.) and injected with either saline (0.9%, s.c.) or semaglutide (0.1 mg/kg, s.c.) 30 minutes prior to their session start. A Two-Way ANOVA of breakpoint across fentanyl doses confirmed significant mean effects of Treatment (F(1,56)=40.48; p<0.05) and Dose (F(3,56)=3.863; p<0.05). Tukey’s post-hoc analysis revealed significant decreases in breakpoint for animals receiving semaglutide versus saline across all tested doses of saline and fentanyl (p<0.05) as well as for breakpoint of saline animals receiving 2.5 and 10 µg/kg/inf (i.v.) compared to those self-administering saline (0.9%, i.v.; p<0.05) as shown in **Figure 3C**. Another Two-Way ANOVA of rewards earned by LgA animals during PR confirmed significant main effects of Treatment (F (1,56)=47.83; p<0.05) and Dose (F(3,56)=3.045; p<0.05). Tukey’s post-hoc analysis revealed significant decreases in breakpoint for animals receiving semaglutide (0.1 mg/kg, s.c.) compared to saline (0.9%, s.c.) across all tested doses of saline and fentanyl (p<0.05; **Figure 3D**).

Following the PR sessions and ∼60 hour abstinence period where animals did not receive fentanyl or semaglutide, the semaglutide-experienced LgA animals completed a dose-response assessment where they received escalating doses of saline (0.9%, s.c.) and semaglutide (0.01-0.1 mg/kg s.c.) while self-administering a 2.5 µg/kg/inf (i.v.) dose of fentanyl under an FR1 schedule of reinforcement. Shown in **Figure 3E**, an ordinary One-Way ANOVA confirmed a significant difference for Treatment with multiple comparisons revealing a significant decrease in rewards earned by animals receiving 0.03 mg/kg semaglutide (s.c.) versus saline (s.c.; p<0.05), 0.1 mg/kg semaglutide (s.c.) versus saline (s.c.; p<0.05), and 0.1 mg/kg semaglutide (s.c.) versus 0.01 mg/kg semaglutide (s.c.; p<0.05).

### 3.5 Semaglutide decreased spontaneous locomotor activity following fentanyl discontinuation

Spontaneous locomotor activity was assessed by examining ambulatory activity, horizontal activity, vertical activity, total distance travelled, and rest time. A Two-Way ANOVA was used to confirm significant effects of ambulatory activity, horizontal activity, vertical activity, total distance travelled, and total rest time. For ambulatory activity (**Figure 4A**), we found significant effects of Treatment (F(1,28)=5.148; p<0.05) and Access Duration x Treatment (F(1,28)=5.757; p<0.05). Tukey’s post-hoc analysis confirmed a significant decrease in ambulatory activity between ShA animals receiving semaglutide (0.1 mg/kg, s.c.) versus vehicle (0.9% saline, s.c.; p<0.05). For horizontal activity (**Figure 4B**), we found significant effects of Treatment (F(1,28)=5.450; p<0.05) and Access Duration x Treatment (F(1,28)=5.701; p<0.05). Tukey’s post-hoc analysis confirmed a significant decrease in horizontal activity between ShA animals receiving semaglutide (0.1 mg/kg, s.c.) versus vehicle (0.9% saline, s.c.; p<0.05). For vertical activity (**Figure 4C**), we confirmed significant effects of Treatment (F(1,28)=7.226; p<0.05) and Access Duration x Treatment (F(1,28)=4.215; p<0.05). Tukey’s post-hoc analysis confirmed a significant decrease in vertical activity between ShA animals receiving semaglutide (0.1 mg/kg, s.c.) versus vehicle (0.9% saline, s.c.; p<0.05). For total distanced travelled (**Figure 4D**), we confirmed significant effects of Treatment (F(1,28)=6.103; p<0.05) and Access Duration x Treatment (F(1,28)=6.927; p<0.05). Tukey’s post-hoc analysis showed a significant decrease in total distance traveled by animals receiving semaglutide (0.1 mg/kg, s.c.) versus vehicle (0.9% saline, s.c.; p<0.05). For total rest time (**Figure 4E**), we found no significant effects of Access Duration (F(1,28)=0.01648; p=0.898756), Treatment (F(1,28)=2.557; p=0.121031), nor Access Duration x Treatment (F(1,28)=3.882; p=0.058763).

**Figure 4.**
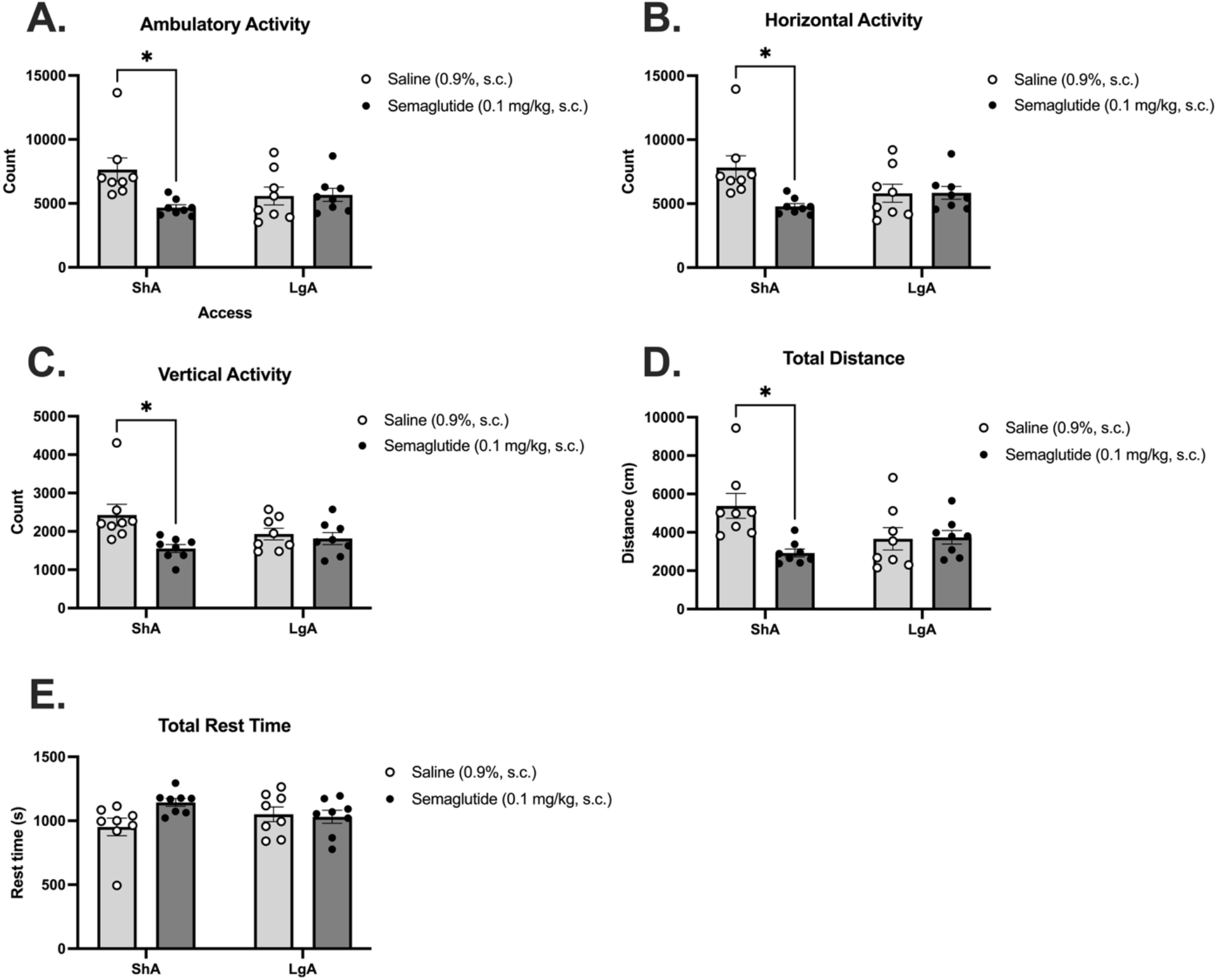
**(A)** Mean ambulatory activity (count ±SEM) for ShA and LgA animals receiving saline or semaglutide 30 minutes prior to assessment. * indicates revealed a significant decrease in ambulatory activity count for ShA animals receiving semaglutide versus vehicle (p<0.05). **(B)** Horizontal activity count for ShA and LgA animals receiving saline or semaglutide 30 minutes prior to assessment. * indicates a significant decrease in horizontal activity count for ShA animals receiving semaglutide versus vehicle (p<0.05). **(C)** Vertical activity count for ShA and LgA animals receiving saline or semaglutide 30 minutes prior to assessment. * indicates a significant decrease in vertical activity count for ShA animals receiving semaglutide versus vehicle (p<0.05). **(D)** Total distance travelled (cm) by ShA and LgA animals receiving either saline or semaglutide 30 minutes prior to assessment. * indicates a significant decrease in total distance travelled by ShA animals receiving semaglutide versus saline (p<0.05). **(E)** Total rest time (s) of ShA and LgA animals receiving either saline or semaglutide 30 minutes prior to assessment. No significant main effects were observed.

### 3.6 Semaglutide did not alter thigmotaxis following fentanyl discontinuation

Two-Way ANOVA was used to confirm significant effects for center duration and perimeter duration during an open field assessment. For center duration (**Figure 5A)**, we found no significant effects of Access Duration (F(1,28)=1.652; p=0.209205), Treatment (F(1,28)=1.648; p=0.209790), nor Access Duration x Treatment (F(1,28)=0.4833; p=0.492645). For perimeter duration (**Figure 5B**), we also found no significant effects of Access Duration (F(1,28)=1.652; p=0.209204), Treatment (F(1,28)=1.647; p=0.209844), nor Access Duration x Treatment (F(1,28)=0.4833; p=0.492643). Our findings indicate that semaglutide dose not significantly alter thigmotaxis.

**Figure 5.**
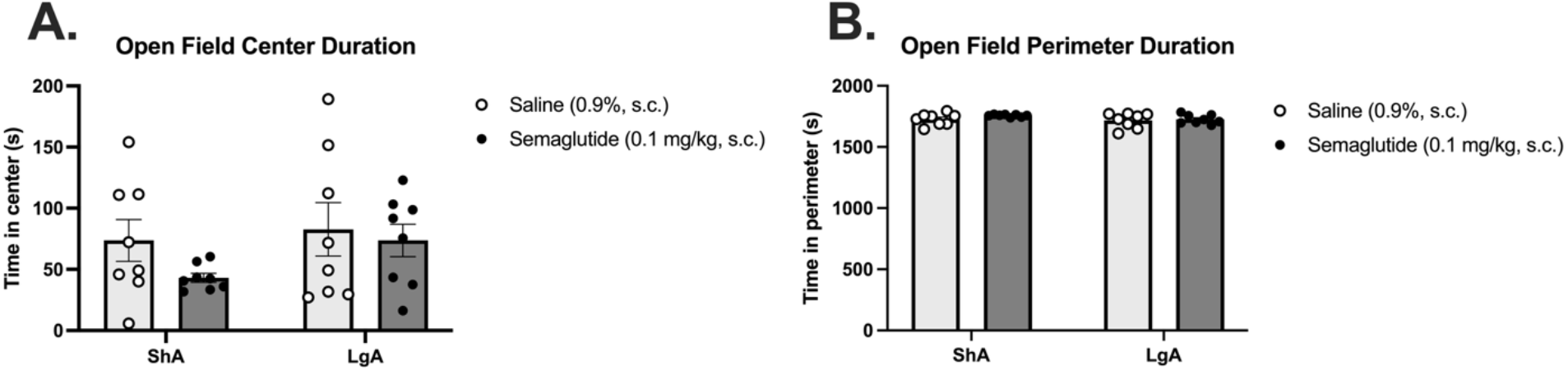
**(A)** Total time in center (seconds ± SEM) during open field assessment for ShA and LgA animals receiving either saline or semaglutide. No significant main effects were observed. **(B)** Total time in perimeter (seconds ± SEM) during open field assessment for ShA and LgA animals receiving either saline or semaglutide. No significant main effects were observed.

### 3.8 Semaglutide decreased opioid withdrawal symptoms in animals trained under ShA conditions

A Two-Way ANOVA of withdrawal scores found a significant main effect of Treatment (F(1,28)=13.70; p<0.05). As shown in **Figure 6**, Tukey’s post-hoc analysis showed a significant decrease in withdrawal scores for ShA animals receiving semaglutide (0.1 mg/kg, s.c.) versus saline (0.9%, s.c.; p<0.05). These findings indicate that semaglutide decreases withdrawal-like behavior in ShA but not LgA animals.

**Figure 6.**
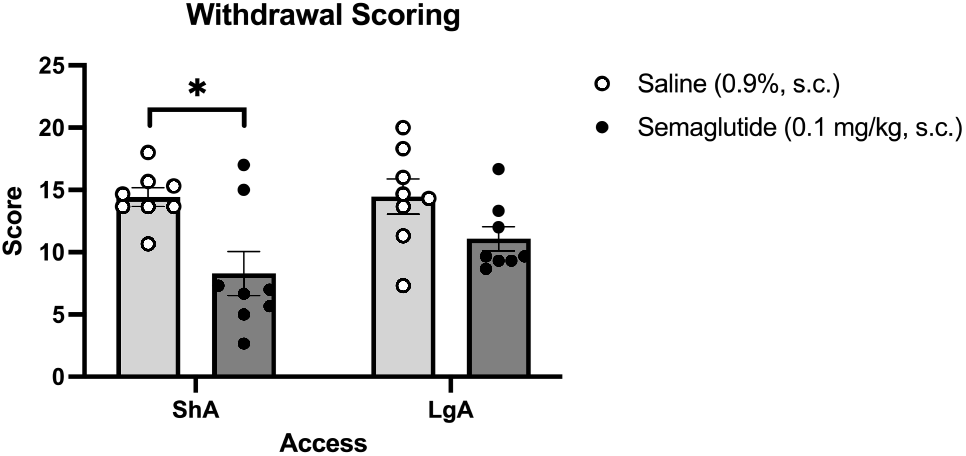
Withdrawal scores (mean ± SEM) for ShA and LgA animals following drug discontinuation. * indicates a significant decrease in withdrawal scores for ShA animals receiving semaglutide compared to saline (p<0.05).

### 3.8 Semaglutide decreased body weight in ShA and LgA rats

A Three-Way ANOVA examining subject body weights throughout the study revealed significant main effects of Session (F(31.00,434.0)=70.86; p<0.05), Session x Access Duration (F(3.352,46.93)=4.859; p<0.05), Session x Treatment (F(31.00,434.0)=36.51; p<0.05), and Session x Access Duration x Treatment (F(3.352,46.93)=2.731; p<0.05). Shown in **Figure 7**, Tukey’s post-hoc analysis confirmed a significant decrease in body weight for ShA animals receiving semaglutide compared to session 21 (the last session before they began receiving semaglutide) from PR session 2 through the first abstinence day (p<0.05) and a significant decrease in body weight for LgA animals receiving semaglutide compared to session 21 from PR session 2 through the first abstinence day (p<0.05).

**Figure 7.**
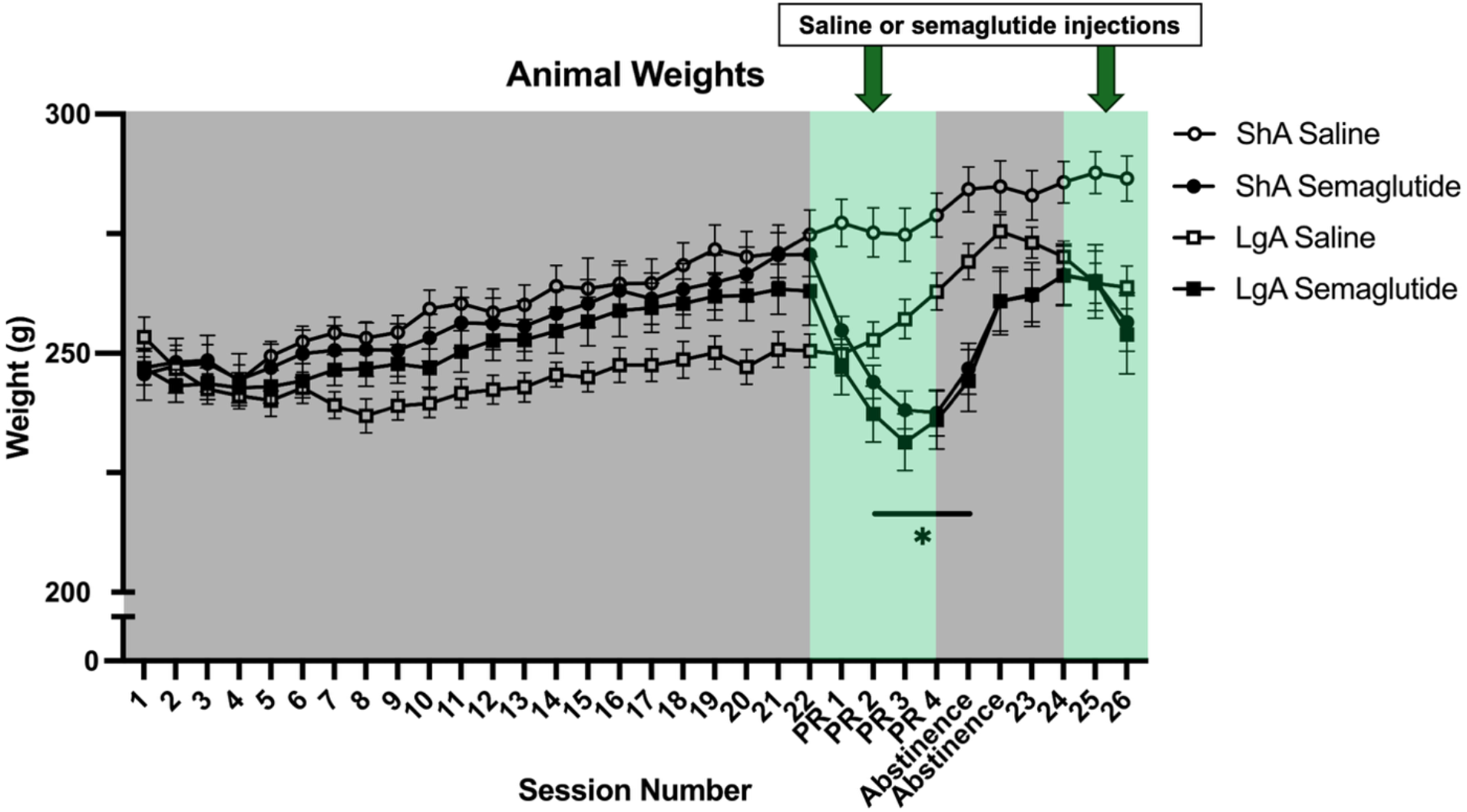
Animal body weight (grams ± SEM) throughout duration of the study. Numbers alone indicate FR1 sessions, blank spaces indicate abstinence days, and PR sessions. Animal weights were recorded prior to semaglutide injections each day. * indicates significantly decreased body weights for ShA and LgA animals receiving semaglutide compared to session 21 (p<0.05).

## 4. DISCUSSION

We found main effects of semaglutide being able to decrease breakpoint under a progressive ratio schedule of reinforcement. These findings, paired with the previously mentioned findings that semaglutide does not alter lever discrimination, show that semaglutide specifically decreases motivation to self-administer intravenous fentanyl. These findings are consistent with previous studies that found other GLP-1R agonists, such as GEP12, can acutely decrease motivation for fentanyl under a progressive ratio schedule of reinforcement [33].This is of clinical relevance and begs further investigation into whether these effects on motivation and/or cravings for opioids translate to patients suffering from OUD. These findings suggest that semaglutide decreases the rewarding value of fentanyl rather than enhancing it, unlike what was observed with the ability of delta(9)-tetrahydrocannabinol to decrease oxycodone self-administration [44]. Previous studies have tended to focus on examining the effects of GLP-1 receptor agonists during drug reinstatement. To the best of our knowledge, the present study represents the first to specifically investigate the effects of GLP-1R agonists on motivation to self-administer drug immediately following acquisition. However, previous studies have examined the ability of GLP-1 receptor agonists to decrease opioid self-administration under a PR schedule of reinforcement following reinstatement, and these studies examining animal subjects under a PR schedule of reinforcement following extinction training and subsequent reinstatement have found that GLP-1 receptor agonists consistently decrease self-administration [30-32,35-38]. These findings are consistent with our findings that semaglutide decreased breakpoint, interpreted as motivation to self-administer fentanyl.

Following administration 30 minutes prior to the start of IVSA sessions, we saw that semaglutide decreased rewards earned by animals trained under long-access conditions and that it did so dose-dependently and without altering lever-discrimination. The dose-dependent effects of other GLP-1R agonists has been recorded previously in other studies examining fentanyl and alcohol self-administration [33,45]. Our finding that semaglutide attenuates fentanyl self-administration under a FR1 schedule of reinforcement in long-access animals is consistent with the study by Douton, et al. ^30^ which found that liraglutide, another GLP-1 receptor agonist, decreased heroin, another opioid, self-administration under long-access (in their case 6 hours) conditions. To our knowledge, this is also the first study to directly examine the effects of semaglutide on lever discrimination. Our study demonstrates that systemic semaglutide does not alter lever discrimination, suggesting it does not interfere with the recall of drug memories.

Intravenous self-administration (IVSA) is a commonly used model of addiction [46]. More recent studies have focused on extended-access conditions, where animals typically have access to drug for four or more hours, which model the compulsive behaviors associated with OUD, and substance use disorder (SUD) more broadly, as well as the escalation of drug use observed in individuals with SUD [39,47]. Additionally, fentanyl dosing in this was consistent with previous intravenous fentanyl self-administration studies, including studies examining various GLP-1R agonists and their effects on fentanyl use [33,36,38]. A study found that GLP-1 receptor agonists can attenuate escalation of drug taking in animals trained to self-administer heroin [30]. Future work should further examine the effects of GLP-1 receptor agonists, such as semaglutide, on escalation during acquisition of opioid self-administration. We found that animals trained under long-access conditions acquired fentanyl IVSA behavior slightly earlier than animals trained under short-access conditions. Additionally, we found that female rats trained under long-access conditions escalated their fentanyl intake, while animals trained under short-access conditions did not. These findings are consistent with previous studies finding that animals trained under extended-access (i.e. long-access) conditions escalate responding and intake while those trained under short-access conditions do not [39,47-49]. Previous work has also shown that female rats escalate drug-taking faster than males [47,48].

To our knowledge, no studies have examined the effects of GLP-1 receptor agonists on opioid withdrawal. We found that semaglutide decreased symptoms of opioid withdrawal in animals trained under short-access but not long-access conditions. Our opioid withdrawal scoring system was adapted from Gellert and Holtzman ^43^; however, it is important to note that their study examined on precipitated withdrawal from morphine following naloxone exposure, whereas the present study observes opioid withdrawal symptoms from fentanyl following spontaneous withdrawal. Furthermore, this study did not use fentanyl-naïve control animals. Therefore, there are some limitations in our interpretation. Our findings suggest that semaglutide does not decrease withdrawal-like symptoms in escalated animals: those trained under long-access conditions and that have a dependence-like phenotype more similar to that of humans with OUD. These findings have great translational relevance and suggest that, as a potential treatment for opioid use disorder, the effects of semaglutide on symptoms of opioid withdrawal may be of lesser benefit to patients with more severe OUD compared to mild or moderate. Another caveat to our findings on opioid withdrawal is that semaglutide appears to have had no effect on anxiety-like behavior. While open field test is often used to assess locomotor behavior, it can also be used to assess anxiety-like behavior in female animals [50]. Our findings regarding anxiety-like behavior are consistent with a different study examining the effects of semaglutide in a model of Alzheimer’s disease, where, similarly, semaglutide had no effect on anxiety-like behavior [51]. Although, our finding that semaglutide can decrease withdrawal-like symptoms in animals trained under short-access conditions is similar to other findings that liraglutide, another GLP-1 receptor agonist, can decrease symptoms of nicotine withdrawal [52]. Additional work is still necessary to understand fully the effects of GLP-1 receptor agonists on opioid withdrawal.

We found that semaglutide decreased spontaneous locomotor activity in animals trained under short-access conditions but not those trained under long-access conditions. This decrease in spontaneous locomotor activity, at least for the animals trained under short-access conditions, is consistent with previous findings that GLP-1 receptor agonists can decrease locomotor activity [23]. Perhaps these findings suggest that more “dependent” animals are already experiencing decreased locomotion, so the addition of semaglutide does not significantly alter locomotion in long-access animals. However, a different study examining the effects of semaglutide on locomotor activity in drug-naïve animals found that semaglutide increased locomotion in both male and female rats [20], a finding that differs from our study and others. Interestingly, another study found that the GLP-1 receptor agonist liraglutide did not alter locomotor activity assessed via the rotarod test [28]. However, it is possible that our findings highlight potential rate-suppressing effects of semaglutide. This possibility is further understood by the observation that semaglutide decreases self-administration of saline under a PR schedule of reinforcement in addition to fentanyl. Although, this effect may also be explained by extinction bursting as the session in which animals received saline (i.v.) was also the first session in which the animals did not receive fentanyl (i.v.).

We confirmed that semaglutide was able to decrease body weight over the course of treatment, and expected outcome of repeated GLP-1 receptor agonist injection. These findings are consistent with previous studies examining the effects of GLP-1 receptor agonism on drug-taking in pre-clinical models [29,32,33,36]. Although Douton, et al. ^30^ found that liraglutide did not significantly impact weight gain in animals trained to self-administer heroin. Interestingly, we also saw that cessation of semaglutide injections caused the animals’ body weight to rebound. Given that research with GLP-1 receptor antagonists has been relatively limited, future investigations could focus on whether administration of opioids also rebound after discontinuation of GLP-1 receptor agonists following chronic use. A study by Marty and colleagues [41] confirmed that semaglutide significantly decreased ethanol intake and found that GLP-1 receptor antagonism by exendin 9-39 did not reverse the suppressing effects of semaglutide in male rats. However, an earlier study by Shirazi and colleagues [53] found that GLP-1 receptor antagonism by exendin-3(9-39) actually increased ethanol drinking also in male rats. As both of these studies were performed in male Wistar rats, future studies should examine the effects of GLP-1 receptor antagonism in female subjects.

Furthermore, due to the conflicting nature of previous studies, future work should elucidate how GLP-1 receptor antagonism may or may not increase drug intake in alcohol and other substances of abuse, such as opioids. Overall, our study addresses a gap in the existing literature by examining the ability of semaglutide to attenuate motivation for opioid self-administration in female subjects. Previous studies regarding GLP-1 receptor agonists and animal models of substance use disorder have tended to focus on male subjects despite over 59% of the 871,854 people prescribed GLP-1 receptor agonists from 2011-2023 being women [16]. Although this study did not address sex as a biological variable, one study found that female rats escalate drug taking faster than male rats [47]. Future investigation could determine whether the effects of semaglutide and other GLP-1 receptor agonists on drug self-administration are sex-dependent. Collectively, these findings suggest semaglutide may modulate motivation to seek opioid reward and could be useful in the development of pharmacotherapies to address OUD.

## 6. AUTHOR CONTRIBUTIONS

**Kaitlyn E. Rojas**: Conceptualization, Methodology, Formal Analysis, Investigation, Data Curation, Writing – Original Draft, Writing – Review & Editing, Visualization, Project administration. **Sarah C. Gee**: Investigation, Writing – Review & Editing. **Chris L. Wernette**: Investigation, Writing – Review & Editing. **Eric X. Wang**: Investigation, Writing – Review & Editing. **Emily T. Nguyen**: Investigation, Writing – Review & Editing. **Jacques D. Nguyen**: Conceptualization, Methodology, Formal Analysis, Writing – Original Draft, Writing – Review & Editing, Supervision, Funding acquisition. All authors approved the final version of this manuscript.

## 7. ACKNOWLEDGEMENTS

Support for this study was provided by NIH DA047413 (JDN) and the Ronald E. McNair Scholars Program Research Fund (KER). The funding bodies had no influence on the study design, data interpretation, manuscript creation or publishing decisions.

## 8. DISCLAIMERS

The authors have no competing interests to declare.

## 9. ETHICAL APPROVALS

All aspects of this study were approved by the Baylor University Institutional Care and Use Committee (IACUC #1984561).

## 10. DECLARATION OF GENERATIVE AI IN SCIENTIFIC WRITING

The authors declare that AI was not used in any part of the writing process.

## 11. DATA AVAILABILITY STATEMENT

Data will be made available to qualified individuals upon legitimate request.

